# Integration of viral transcriptome sequencing with structure and sequence motifs predicts novel regulatory elements in SARS-CoV-2

**DOI:** 10.1101/2020.06.24.169144

**Authors:** Brian J. Cox

## Abstract

In the last twenty years, three separate coronaviruses have left their typical animal hosts and became human pathogens. An area of research interest is coronavirus transcription regulation that uses an RNA-RNA mediated template-switching mechanism. It is not known how different transcriptional stoichiometries of each viral gene are generated. Analysis of SARS-CoV-2 RNA sequencing data from whole RNA transcriptomes identified TRS dependent and independent transcripts. Integration of transcripts and 5’-UTR sequence motifs identified that the pentaloop and the stem-loop 3 were also located upstream of spliced genes. TRS independent transcripts were detected as likely non-polyadenylated. Additionally, a novel conserved sequence motif was discovered at either end of the TRS independent splice junctions. While similar both SARS viruses generated similar TRS independent transcripts they were more abundant in SARS-CoV-2. TRS independent gene regulation requires investigation to determine its relationship to viral pathogenicity.

## Introduction

While coronaviruses (CoV) are endemic to much of the world, many are relatively harmless, contributing along with other viral species to the common cold in humans(Coleman and Frieman, 2014). However, three recent events of coronaviruses jumping hosts have caused outbreaks of lethal pulmonary infections. The SARS subtype infected thousands of people and killed 800 or nearly 10% of all infected individuals. MERS was more harmful but spread poorly between individuals(Coleman and Frieman, 2014). The most recent event COVID-19 (formerly 2019-nCoV) began late 2019 is currently ongoing with millions infected and 100s of thousands of deaths. Also, the preventative measures to reduce the spread of the virus is causing global economic dysfunction. Intense global efforts are ongoing to develop vaccines and drug treatments. An assessment of the regulation and expression of viral genes is needed to assist in the development of diagnostics, vaccines and pharmaceutical interventions.

The coronavirus is a family of positive single-stranded RNA viruses that use only RNA to generate RNA copies of its genome and subgenomic messenger RNA (sgmRNA) transcripts of specific coding sequences(Sola et al., 2011; Yang and Leibowitz, 2015). RNA folding is essential to the function of RNAs, and the CoV is no exception, with many predicted and verified stem-loop (SL) structures and sequence motifs essential to the viral life cycle(Sola et al., 2011, 2015). After infection, the viral genome is loaded on to the ribosome, and the first open reading frame ORF1a is transcribed as a polycistronic message(von Brunn et al., 2007; Sola et al., 2011). The polypeptide is cleaved by an encoded protease to release 15-16 non-structural proteins involved in RNA replication. Another more extended polypeptide is expressed by a slip sequence consisting of two stem-loops at the end of Orf1a. Within the ribosome, the slip sequence shifts the reading frame to translate the Orf1b sequence.

Downstream of Orf1ab are 9-10 open reading frames that become expressed through the formation of specific transcripts called subgenomic messenger RNAs (sgmRNA)(Sola et al., 2015). Essential to sgmRNA synthesis are conserved sequences called the TRS-L in the 5’-UTR and the TRS-B located upstream of multiple protein-coding genes. sgmRNAs of the CoV genome begin in the 3’UTR where the viral RNA directed RNA polymerase (Rdrp) and its associated proteins create a negative RNA copy to the viral genome(Kaye et al., 2006; Sola et al., 2015; Yang and Leibowitz, 2015). During the viral replication process, the Rdrp can skip the intervening sequence and bridge a TRS-B sequence to the TRS-L of the 5’UTR, termed template switching(Sola et al., 2015). The negative transcripts are next transcribed into positive strands and loaded on to the ribosome for the production of viral proteins. The sgmRNA transcripts are differentially expressed by unknown mechanisms to maintain a stoichiometric balance of transcripts for protein production and viral assembly.

The TRS-L is part of a series of folded structures within the UTRs and is essential to both viral genome replication and transcription(Yang and Leibowitz, 2015). The folds of the 5’ and 3’ UTRs RNA of the coronavirus family are modelled using thermodynamic systems to find the lowest free energy state. The models are compared to the solved structure of betacoronavirus 1 (more commonly known as bovine coronavirus, BCoV) and murine coronavirus (MHV). In studies on the SARS and MERS viruses, conserved SL structures were observed(Sola et al., 2011). However, few studies have attempted to discover these sgmRNAs in CoV directly. Only one recent study examined the expression of sgmRNAs during SARS-CoV-2 infection(Kim et al., 2020). They observed that >95% of the viral transcripts used the TRS dependent mechanism. The TRS independent sgmRNAs may be largely non-coding and showed increased presence of RNA methylation. However, their methods for detecting and quantifying sgmRNAs used poly-A dependent sequencing mechanisms, leaving the possibility that non-polyadenylated sgmRNAs exist.

This current study aims to identify possible novel mechanisms of sgmRNA regulation that may be useful in developing immunization strategies, therapeutic drugs and improving our understanding of CoV biology. I assessed the 5’ and 3’UTRs using structure prediction methods to characterize the similarities of the folded structures and locations of the TRS to other CoV. Secondly, a detailed analysis of RNA-sequencing data from SARS-CoV-2 and SARS-CoV infected cell and animal models was conducted to determine the locations of spliced sgmRNAs and their relative abundance. Sequence motif discovery and analysis were conducted with structural prediction to assess the relationship to sgmRNA abundance. TRS independent transcripts were assessed to identify possible known and novel regulatory motifs.

## Results

### SARS-CoV-2 has conserved 5’ and 3’ UTR structural elements with other CoV

Based on previously published analysis of the 5’- and 3’-UTR regions of several CoV family members, there are conserved sequence motifs and folded RNA structures essential to the regulation of viral genome replication and sub-genomic mRNA (sgmRNA) production(Sola et al., 2011). I assessed the sequence and structural conservation of the 5’UTR regions of SARS-CoV-2 against SARS-CoV and MERS (**Figure 1**). In all three genomes, biomolecular prediction of the first 130 bases predicted four conserved stem-loops (SL) were observed: SL1 (involved in genomic replication(Zhang et al., 1994)), SL2 (a conserved pentaloop sequence typical to many CoV species(Zhang et al., 1994)), SL3 (containing the TRS-L sequence that is essential for the generation of sgmRNA(Zhang et al., 1994)) and SL4. While previously published predictions of the MERS 5’UTR indicated that the SL3 structural element is absent(Sola et al., 2011), I predicted an SL3 stem-loop structure. There is no wet-lab experimental evidence of these stem-loops in SARS-CoV, SARS-CoV-2 or MERS, and their functions are inferred based on related CoV species with experimental evidence.

**Figure 1.**
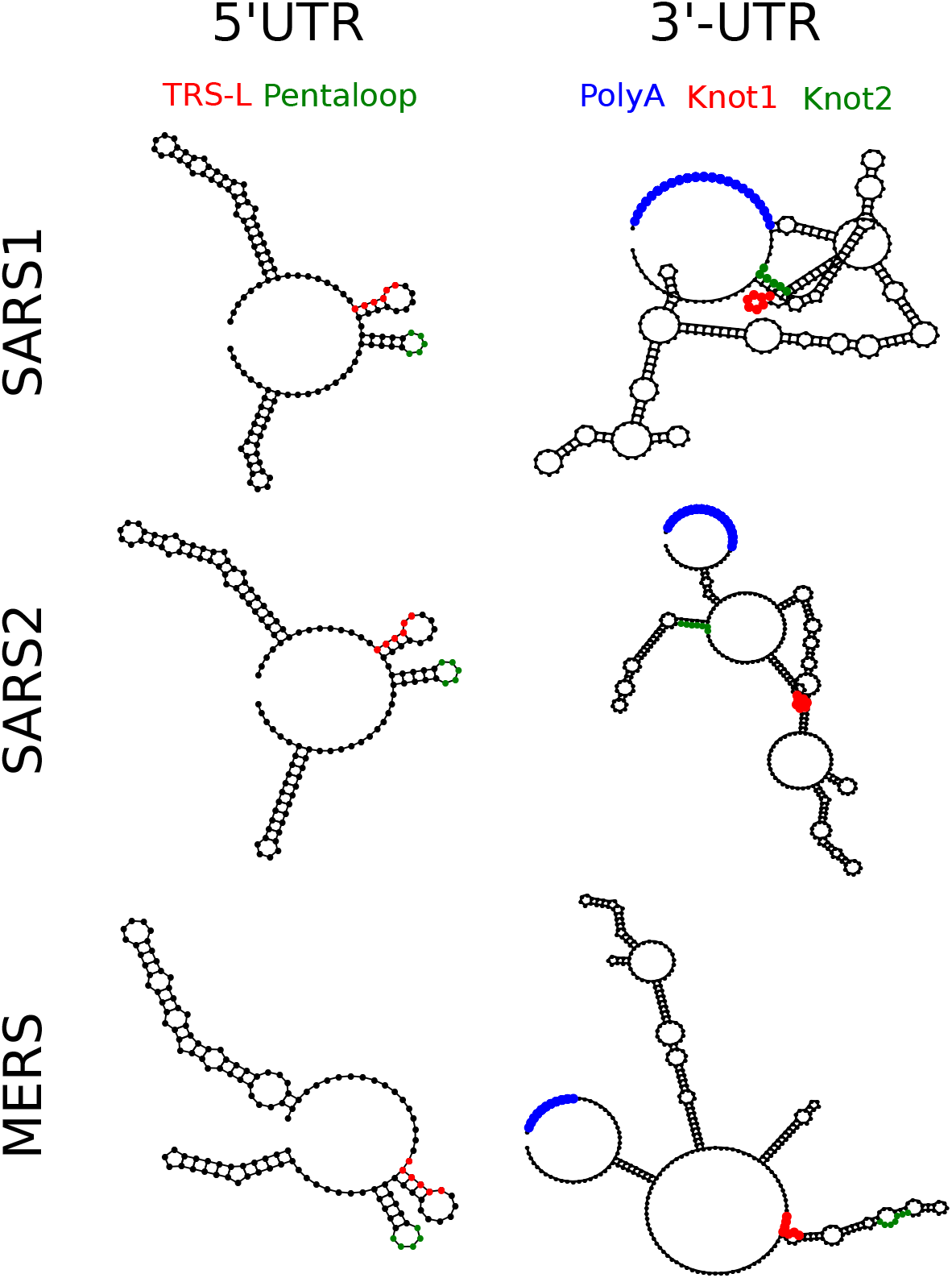
Untranslated regions of human coronaviruses have common folded structures. The first 130 bases of the SARS-CoV, SARS-CoV-2 and MERS genomes were folded using RNAStructure and visualized using the R library *RRNA* function *RNAPlot*. The TRS-L and the pentaloop region of SL2 are highlighted. In all cases, the regulatory TRS and pentaloop are on two adjacent stem-loops, with the TRS located in the stem portion. A similar analysis of 350 bases of the 3’UTR essential for viral replication and negative-strand transcript initiation shows more diversity in predicted structures. The pseudoknot sequences are conserved in red and green. Additionally, in all species, the poly-A region (blue) is predicted to be looped back to 5 prime regions thought to be important in self-priming the viral RNA directed RNA polymerase.

### SARS-CoV-2 subgenomic RNAs

Given the presence of conserved 5’UTR sequences and structures in SARS-CoV-2, including the TRS-L, sgmRNA should be observed in RNA sequencing data of infected cell or animal models. A recently published study provided evidence of TRS dependent as well as independent sgmRNAs of the SARS-CoV-2 genome(Kim et al., 2020). They reported that TRS independent sgmRNAs represented only a minority of the total transcript (1.5%). However, the sequencing methods utilized in that study could only capture RNAs with a polyadenylation sequence. It is not known if all sgmRNA are polyadenylated. Some RNAs, especially non-coding RNAs, generally lack a poly-A tail, which could explain poor detection of TRS independent sgmRNAs.

Using published sequencing data of ribosome depleted total RNA from SARS-CoV-2 infected cells and animals(Blanco-Melo et al., 2020), I aligned these against the viral genome (**Figure 2**). Using the aligned reads, I generated transcript models using *stringtie*, which identified multiple spliced species that aligned with the TRS-L templated events (**Figure 2**). TRS-L transcripts aligned to the genes S, ORF3a, ORF7a and N. The S transcript also presented with a short deletion of 36 bases.

**Figure 2.**
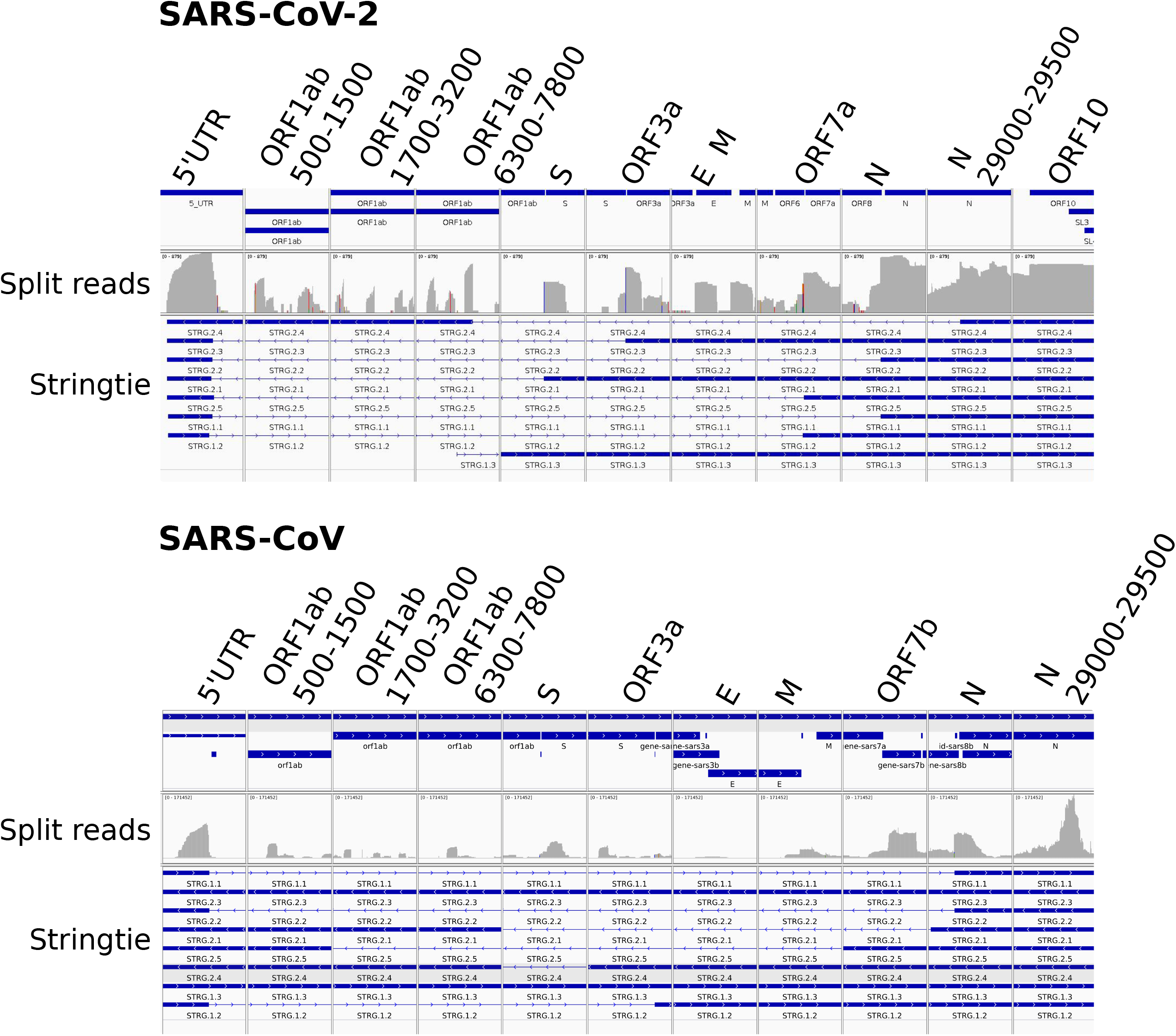
*Stringtie* predicts SARS-CoV and SARS-CoV-2 TRS dependent and independent transcripts. Integrated genome viewer transcripts are shown. For both species, the top track is the gene annotation, the middle track is the split read alignment histograms for regions with peaks of aligned reads (presented as a log scale), and the third track is stringtie predicted transcripts. Peaks of alignment are observed for the 5’UTR leader, internal to the ORF1ab genes and the beginning of S, ORF3a, E, M ORF7a and N as well as internal to N. Bottom track is the predicted transcript from *stringtie*. Both SARS viruses have ORF1ab split reads aligning to ORF1ab and to the middle of the N gene that does not conform to TRS dependent mechanisms of sgmRNA generation.

Also, TRS independent spliced transcripts were identified. Non-canonical transcripts spliced from ORF1ab 7016 onto ORF1ab 19560 and from ORF1ab 7278 onto N at base 29189. The dominant ORF1-N fusion product (bases 7,278 to 29,189) is predicted to have an open reading frame for part of ORF1ab that includes the leader protein (nsp1), nsp2 and approximately 77% of nsp3. For nsp3, this would create the near full-length protein, including the nucleic acid-binding domain (NAR), but would remove the transmembrane domain. The fused N gene is out-of-frame and would not create a protein product. This same ORF1ab-N fusion was previously reported by one other group as one of the non-canonical sgmRNA species in SARS-CoV-2(Kim et al., 2020).

The *stringtie* quantification of the predicted spliced products showed that relative transcripts abundance of TRS dependent was 92.7% while the TRS independent was 7.3%. Of the TRS independent sgmRNAs, the ORF1ab-N spliced product represented 6.6% of the predicted spliced RNA, which was the fourth most abundant of the 8 predicted sgmRNA species, more than several of the TRS-dependent sgmRNAs. While this finding represents an approximately four times higher detection of TRS independent sgmRNAs than previously reported, the methods of quantification in these two studies are not directly comparable, challenging the determination of the relative abundance. Another caveat is that *stringtie* uses mammalian splicing rules that would not follow in the splicing of viral RNA that uses a template switch process.

To better identify possible sgmRNA that may be independent of poly-A, split reads aligned to the viral genome were extracted and quantified (**Table 1**). Using this metric, TRS dependent sgmRNAs that spliced the 5’TRS-L onto TRS-B sequences upstream of viral genes represented only 56% of the reads and matched the genes S, ORF3a, E, M, ORF7a, ORF8 and N, identifying multiple TRS dependent spliced products missed by *stringtie*. Moreover, TRS independent reads were identified with multiple sgmRNSAs mapping from six different regions of ORF1ab into two regions of the N gene (**Table 1**).

**Table 1.** Split reads for SARS-CoV-2 mapping locations and depth

| <u>Split 5'</u> |  |  | <u>Split 3'</u> |  |  | Read<br>Depth | Fraction<br>of all<br>counts | strand |
| --- | --- | --- | --- | --- | --- | --- | --- | --- |
| Start | End | Name | Start | End | Name |  |  |  |
| 13 | 63 | 5'UTR | 21549 | 21682 | S | 39 | 0.03 | - |
| 13 | 65 | 5'UTR | 25380 | 25513 | ORF3a | 172 | 0.11 | - |
| 28 | 69 | 5'UTR | 26236 | 26377 | E | 35 | 0.02 | - |
| 10 | 64 | 5'UTR | 26467 | 26607 | M | 26 | 0.02 | - |
| 10 | 60 | 5'UTR | 27378 | 27499 | ORF7a | 17 | 0.01 | - |
| 10 | 66 | 5'UTR | 27384 | 27519 | ORF7a | 66 | 0.04 | - |
| 16 | 65 | 5'UTR | 27883 | 28005 | ORF8 | 24 | 0.02 | - |
| 11 | 64 | 5'UTR | 28254 | 28382 | N | 445 | 0.29 | - |
| 11 | 64 | 5'UTR | 28254 | 28380 | N | 33 | 0.02 | + |
| 2824 | 2956 | ORF1ab | 28346 | 28459 | N int | 15 | 0.01 | - |
| 6492 | 6638 | ORF1ab | 28492 | 28587 | N int | 14 | 0.01 | - |
| 603 | 614 | ORF1ab | 29144 | 29279 | N int | 12 | 0.01 | - |
| 7136 | 7278 | ORF1ab | 29188 | 29300 | N int | 264 | 0.17 | - |
| 1863 | 1974 | ORF1ab | 29239 | 29376 | N int | 34 | 0.02 | - |
| 6518 | 6607 | ORF1ab | 29296 | 29425 | N int | 41 | 0.03 | - |
| 597 | 734 | ORF1ab | 29352 | 29431 | N int | 50 | 0.03 | - |
| 2361 | 2482 | ORF1ab | 29402 | 29546 | N int | 16 | 0.01 | - |
| 23458 | 23593 | S int | 23629 | 23761 | S int | 210 | 0.14 | - |
| 23491 | 23594 | S int | 23615 | 23729 | S int | 21 | 0.01 | - |

To explore if TRS independent transcripts are common in other human CoV pathogens, I assessed SARS-CoV(Josset et al., 2014; Xiong et al., 2014) and MERS (Data: PRJNA233943; publication not known) using previously published polyA independent RNA sequencing data sets that also used ribosome depletion of total RNA. For SARS-CoV, I observed that TRS dependent split junctions represented 63% of all mapped split reads (**Figure 2 and Table 2**). This is similar to my observations of SARS-CoV-2, indicating that TRS independent splicing is abundant and likely not polyadenylated. Significantly the ORF1ab-N fusion products were also observed in SARS-CoV but at lower levels (**Table 2**). In the MERS virus, the TRS independent split read events represented 13.7% of all split reads (data not shown), indicating a higher prevalence of TRS dependent sgmRNAs compared to both of the SARS-CoV.

**Table 2.**
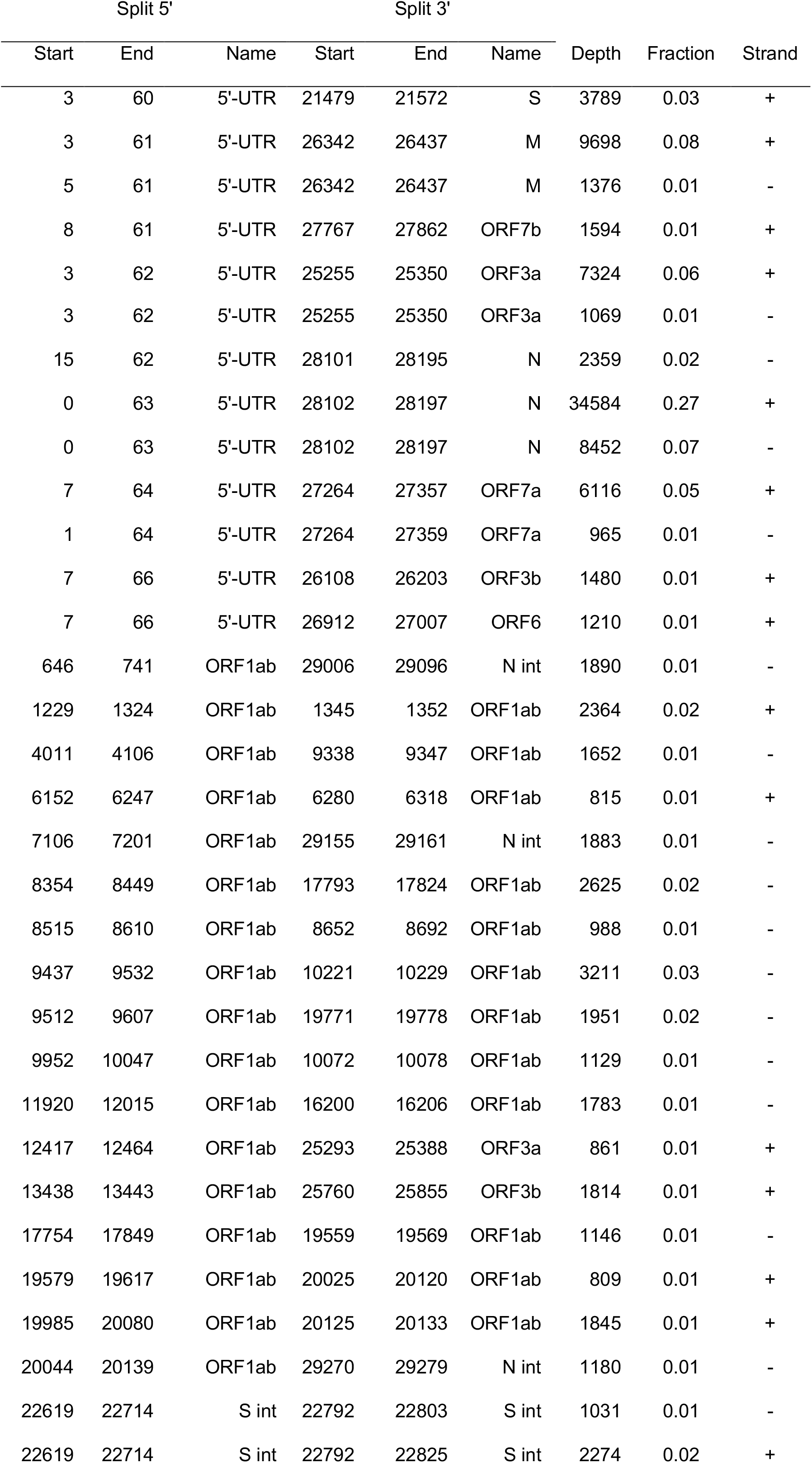
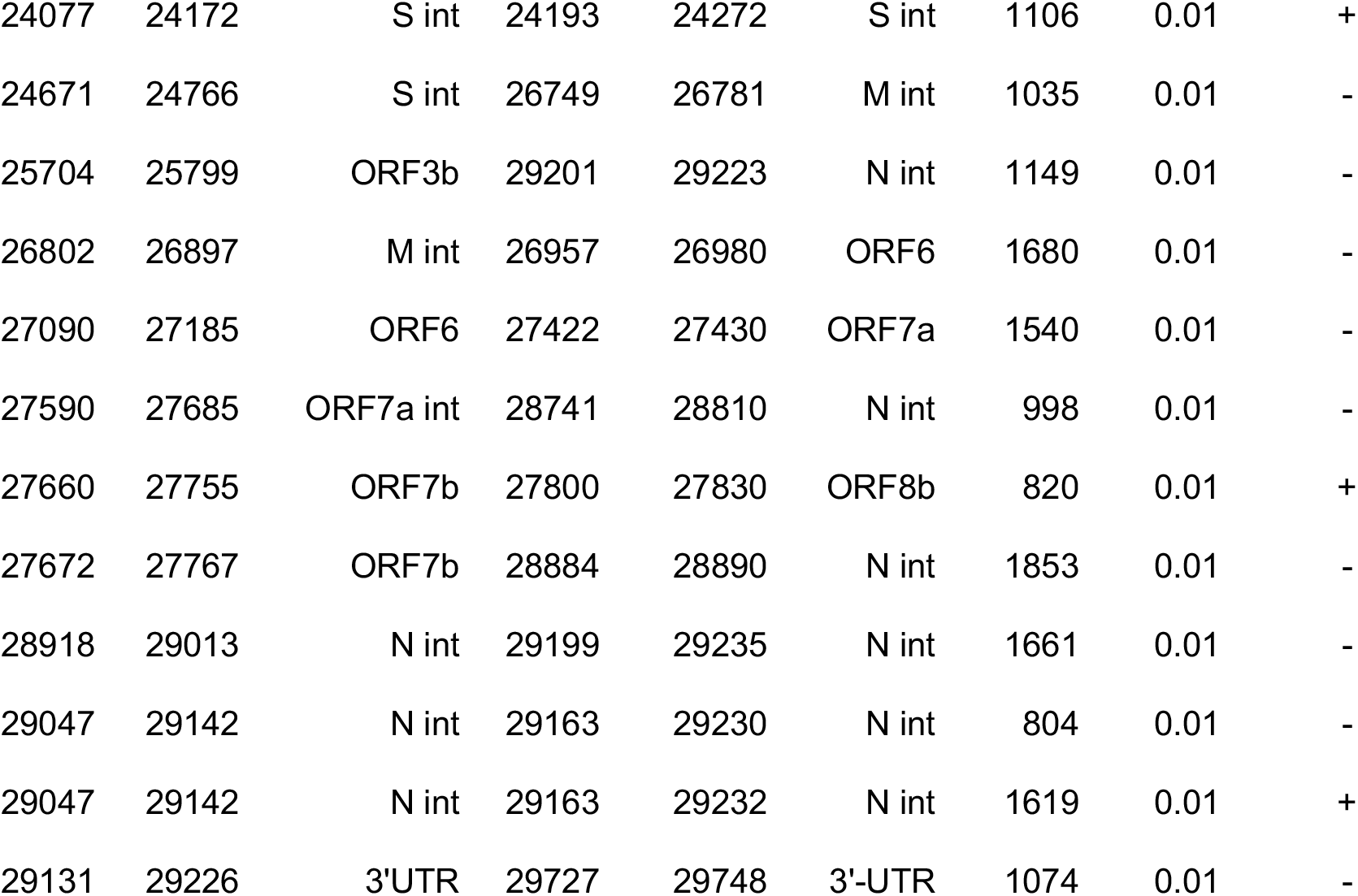
Split reads for SARS-CoV mapping locations and depth

### CoV regulatory sequences for sgmRNA production

To identify those splice variants driven by the TRS, I scanned the SARS-CoV-2 genome for exact matches to the TRS motif (ACGAAC) and displayed matching locations along with the position of coding regions and splice junctions (**Figure 3 A-G**). The TRS motif (**Figure 3 C, D, F and G**; red box) was found upstream of the sgmRNA dependent genes (**Figure 3 B-D**; red arcs) that included the genes S, Orf3a, E, M, Orf7a, Orf8 and N. The TRS sequence was never detected within the ORF1ab gene locus. These findings are similar to previously annotated TRS locations

**Figure 3.**
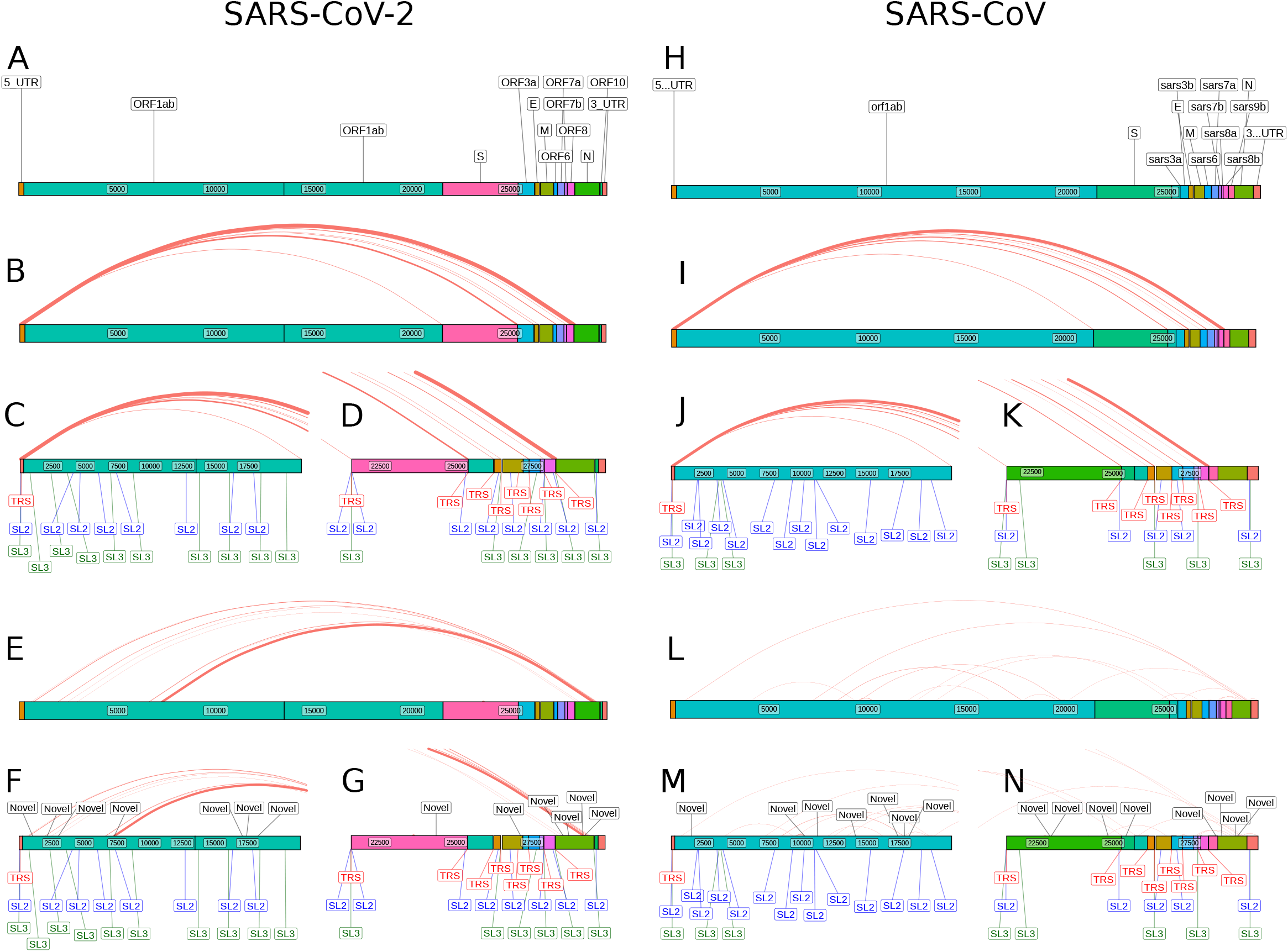
Regulatory sequences drive both TRS dependent and independent sgmRNAs. Scaled images represent the viral genome as coloured bars for each open reading frame and 5 and 3’ untranslated regions. The viral genes are annotated (A and H), and TRS dependent spliced reads are defined as those where the split read maps near the TRS-L and a TRS-B sequence motif (Red boxes). The locations of the regulatory elements of the SL2 pentaloop (Blue boxes) and the SL3 loop (Green boxes) are shown. The spliced reads are represented as red arcs over the whole genome sequence (B and I) with expanded focused views of the 5’ (C and J), or 3’ (D and K) ends of the genomes. TRS independent split reads defined as those that do not map near a TRS signal on both ends of the split reads are displayed as red arcs (E and L) with expanded views of the 5’ (F and M) and 3’ (G and N) genome. The locations of the enriched novel motif found near TRS independent split reads are shown as black boxes (F, G, M and N). All figures are drawn to scale using the R library *ggplot2* graphing functions of *geom_rect, geom_text, geom_curve* and *geom_label_repel*.

Each sgmRNA is expressed at different levels and may be driven by other regulatory elements. As the SL2 conserved loop sequence is also known to be essential for viral sgmRNA, I scanned for this sequence (**Figure 3 C, D, F and G**; blue box). For sgmRNA dependent genes, the SL2 pentaloop sequence (UCUUGU) was discovered only near the 5’ region often near the TRS-B sequence. The pentaloop sequence was also identified in multiple locations within the ORF1ab gene locus. The TRS is located in the stem of SL3, but the loop sequence is also highly conserved in CoV species. For these reasons, I next scanned the SARS-CoV-2 genome for matches SL3 (CUAAAC) loop sequences (**Figure 3 C, D, F and G**; green box). A high correlation was noted in the presence of SL3 sequences near the TRS sequence in the 5’ region of sgmRNA dependent genes. As well, multiple SL3 motifs were observed within the ORF1ab open reading frames. There was no statistical relationship to the presence of SL2 or SL3 sequences and the level of sgmRNA expression (data not shown).

To determine if the TRS, SL2 and SL3 are conserved to other CoV, I repeated the search for these sequences motifs within the SARS-CoV genome (**Figure 3 H-K**). While the TRS and SL2 sequence frequencies and locations were similar between the two SARS viruses (**Figure 3 C, D, J and K**), an exception was that SARS-CoV had half the number of SL3 motifs in the ORF1ab open reading frame compared with SARS-CoV-2 (**Figure 3 C and J**).

While the presence of an RNA sequence motifs was not sufficient to explain sgmRNA abundance, regulatory motifs may need to be located on folded structures such as SL to be active. Next, I assessed the folded structures within the TRS-B loci near sgmRNA of TRS dependent genes to determine if structure and sequence were related to expression levels (**Figure 4**). While a statistical inference is not possible, there was an interesting observation that the TRS-B within the N gene locus folded to place the TRS, SL2 and SL3 sequences on two adjacent stems. N was the only gene with this structure of the TRS-B/SL2/SL3. Only S and E were predicted to have the TRS-B on a stem, but the SL2 and SL3 were unfolded. The SL2 sequence upstream of ORF3a and M was predicted to be on a folded stem, but the associated TRS-B was not. Given that the N gene is the most robustly expressed sgmRNA (4-20 fold higher), the folded structures containing all three sequence elements may be a driver of high template switching.

**Figure 4.**
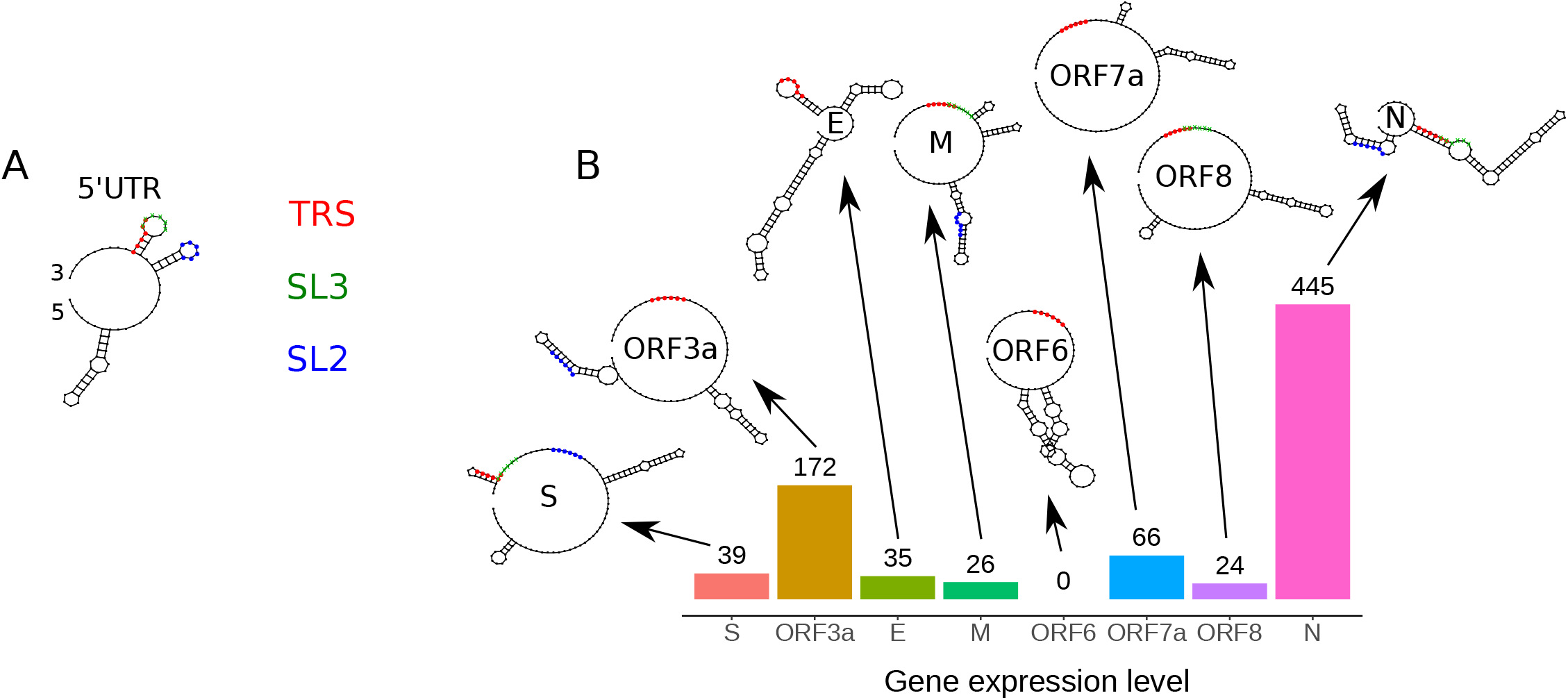
The expression level of the SARS-CoV-2 TRS dependent sgmRNAs related to regulatory sequences and locally folded RNA structures. For comparison (A), the 5’UTR folded structure is shown with the location of the TRS, SL2 and SL3 loops. The local predicted folded RNA structures and the regulatory sequences are displayed with the sequence alignment depth of the split reads that map to each gene. Of note is that only the N gene, with 2.5 to 18 times more expression than all other transcripts, has all regulatory elements predicted to folded stem-loop structures analogous to the 5’ UTR.

A high level of unexpected splicing within the large ORF1ab gene was noted in multiple locations, and all of these formed fusions within the N gene (**Figure 3 E-G**). No TRS was identified within the ORF1ab locus or the N gene near the splice locations. However, within the ORF1ab locus, multiple SL2 and SL3 sequences were identified, but these were not observed within the N gene. An inspection of the ORF1ab splice regions with the N splice regions did identify a novel sequence element (ATTGGC). Searching the SRAS-CoV-2 genome for this element revealed it to be present at 4/6 ORF1ab splice starts, and both N splice ends, including the abundant ORF1ab (base 7,278) to N gene (base 29,189) fusion sgmRNA (**Figure 3 F and G**; black boxes). The novel sequence was also found near SL2 and SL3 motifs in the ORF1ab. Significantly, the novel motif was not observed in the 5’UTR or near TRS-B sequences of genes. I called this motif the TRS independent regulatory (TIR) motif.

Next, I searched for the TIR motif associated with TRS-independent sgmRNAs within the SARS-CoV genome (**Figure 3 L-N**). I observed that the TIR sequence was identified in SARS-CoV at similar locations, as observed in SARS-CoV-2 (**Figure 3 F, G, M and N**). The TIR sequence was found near the splice locations of the in the ORF1ab-N fusion in SARS-CoV (**Figure 3 M**). This suggests a level of conservation for the TIR motif. MERS showed poor conservation of the SL3 motif and the TIR motif (data not shown).

## Discussion

The TRS mechanism of sgmRNA generation is characterized in CoV species other than the two SARS viruses. I show that the folded structure of the 5’-UTR of SARS-CoV-2 is predicted to be similar to SARS-CoV and even MERS. The high conservation indicates that the TRS mechanism is likely used in SARS-CoV-2. A recent report highlighted the presence of TRS independent sgmRNAs and the role that methylation signalling may play in marking these transcripts. In this current work, I draw attention to the possible difference in abundance of the TRS independent sgmRNAs when using total versus poly-A enrichment for library construction. Many RNAs do not contain poly-A sequences, specifically those that function as non-coding RNAs(Zhang et al., 2014). I identified TRS-independent transcripts as more abundant in total RNA compared to poly-A RNA. The lack of poly-A in TRS independent sgmRNAs suggests that they may function as non-coding RNAs. Potential functions of the non-coding RNAs are to inhibit cellular antiviral responses or to amplify the viral genome and sgmRNA by priming RNA-RNA interactions and the viral polymerase.

Methods such as stringtie performed poorly at identifying sgmRNAs, and instead use of split reads gave better performance. Using the depth of reads mapped to the junctions provided quantification of the abundance of TRS dependent and independent reads. Quantification of junctions revealed that the SARS family of CoV had a higher abundance of the TRS independent sgmRNAs compared to MERS.

Viral sgmRNA exist in different proportions that are relative to the protein’s proportion needed for viral particle assembly(Kim et al., 2020; Sola et al., 2015). These different stoichiometries may indicate that there is a higher preference for some template switch regions leading to the increased production of these sgmRNAs over others. To regulate the abundance of sgmRNA transcripts, the TRS likely does not work alone in priming the template switch. To investigate possible combinations of other regulatory motifs, I used the SL2 pentaloop and the SL3 loop upstream of the TRS-L that are part of the 5’-UTR regulatory regions. I noted the presence of the SL2 loop motif upstream of genes that generate sgmRNAs. I also found evidence of an expanded TRS signal that includes the loop of stem-loop 3. While I found no evidence for a correlation of motifs and expression levels, there did appear to be a relationship between the predicted folded structures of the 5’ regions of the viral gene and the regulatory sequences. This relationship was most apparent for the N gene, where the TRS, SL2 and SL3 were all located on two different stems, which was not an arrangement observed on the other viral genes. This folded arrangement may increase the N gene’s affinity for template switching to the 5’-UTR through higher affinity RNA-RNA interactions. Other regulatory RNA elements may be present that explain sgmRNA abundance variation. Alternatively, variation in transcript stability may cause apparent differential expression.

In a comparison of SARS-CoV and SARS-CoV-2, the SL2 and SL3 motifs are preferentially located near TRS-B sequences and not interior to the open reading frames. The exception is ORF1ab, that while lacking TRS motifs, had a high number of SL2 motifs in both species. Interestingly, the SL3 motif frequency was markedly reduced (>50%) within the ORF1ab region of SARS-CoV compared to SARS-CoV-2.

Split reads also identified TRS-independent sgmRNAs. In support of my observations on SARS-CoV-2, I also assessed SARS-CoV and MERs transcriptional data, two other human pathogenic CoV. Significantly, all three of these viruses produce TRS independent transcripts, indicating that this is a broader phenomenon of CoV and not species-specific. I also identified the TIR motif, a novel sequence element (ATTGGC) flanking the spliced regions of TRS independent sgmRNAs in both SARS-CoV-2 and SARS-CoV. The novel sequence element is absent from the 5’UTR and near TRS-B sequences. I observed that while SARS-CoV-2 and SARS-CoV both expressed ORF1ab-N fusions, SARS-CoV-2 had a higher level of expression relative to TRS dependent sgmRNAs. Considering that fewer SL3 motifs were present in SARS-CoV relative to SARS-CoV-2, this suggests that the SARS-CoV-2 virus may be evolving other regulation of sgmRNAs. While SARS-CoV and SARS-CoV-2 are closely related, SARS-CoV-2 has infected more people in more counties. The increased expression of the TRS-independent sgmRNAs may provide a selective advantage to SARS-CoV-2 in human disease relative to SARS-CoV. Mutagenic studies are needed to validate the TIR motif’s role in TRS independent sgmRNA generation.

In conclusion, the novel regulatory sequence and confirmation of TRS independent sgmRNAs will lead to an improved mechanistic understanding of sgmRNA production. The sgmRNA mechanism could be drugged to inhibit viral transcription beyond targeting the polymerase.

## Acknowledgements

This work was funded in part by a Canada Research Chair to B.C. I would like to acknowledge members of my research group for feedback on the analysis and manuscript.

## Author Contribution

B.C. designed and conducted the experiments, analyzed the data and wrote the paper.

## STAR methods

### Viral sequences

All viral genome sequences were obtained from NCBI, SARS-CoV-2 NC_045512.2, SARS-CoV NC_004718.3 and MERS NC_038294.1.

### Alignments and sequence organization

All sequences were handled and processed in R using the biostrings package, including conversion of DNA into RNA, reverse complementation, extraction of subsequences and motif mapping. Related sequences were found using the NCBI BLAST tool(Johnson et al., 2008). Data were visualized using ggplot2(Wickham, 2009).

### RNA folding

RNA folding experiments were performed using the RNA structure web server and stand-alone program(Mathews, 2014) using default conditions.

### RNA-seq data processing

Data sets were obtained from the short read archive (SARS-CoV-2 GSE147507 (Blanco-Melo et al., 2020); SARS-CoV PRJNA227801 (Josset et al., 2014; Xiong et al., 2014); MERS PRJNA233943 (Not linked to a pubmed article)). These included all cell lines, human samples and animal models infected with the virus. FASTQ files were deposited as processed files with adaptors trimmed and poor-quality sequences removed and trimmed. The htseq2(Anders et al., 2014) program was used to generate the viral reference genome and for alignments of the FASTQ files. Reads that did not align with the viral genome were discarded. The setting for alignment included *-data* to prepare for transcript assembly and reducing the non-canonical splice penalty to 0. Aligned reads in sam format were converted into bam files using samtools. Bam files were sorted and indexed using samtools. Reads that split and represented splicing were extracted by using the CIGAR codes by an AWK command to select specific reads that were split. Aligned reads were visualized in the Integrated Genome Viewer.

### De novo transcript identification

Bam files were converted into FASTQ files using the bam2fastq function. Converted FASTQ files were assembled into transcripts using stringtie. The gtf output files from stringtie were visualized in IGV and extracted as a bed file with a 12 column format. The bed12 file was used to extract the stranded fasta sequence from the reference genome. The NBCI ORF finder tool was used to identify open reading frames in novel splice products.

### Code availability

All code is available upon requires

